# *MDNA* : A software module for DNA structure generation and analysis

**DOI:** 10.1101/2025.07.26.666940

**Authors:** Thor van Heesch, Enrico Skoruppa, Peter G. Bolhuis, Helmut Schießel, Jocelyne Vreede

## Abstract

Exploring the dynamical and structural properties of molecular complexes involving DNA is a fundamentally important aspect of understanding many biological processes. Although tools exist for modeling linear DNA and simple complexes, significant challenges remain in generating intricate biomolecular assemblies and incorporating biologically relevant modifications. These limitations restrict the ability to create accurate starting configurations for advanced molecular simulation studies. Here, we introduce `MDNA`, a molecular modeling toolkit that bridges these gaps by enabling the construction and analysis of complex DNA structures. `MDNA` offers a versatile solution to generate DNA shapes using a spline-based mapping technique that enables the construction of DNA configurations with arbitrary shapes. Key features include support for (non-)canonical base modifications, such as Watson-Crick-Franklin to Hoogsteen transitions, DNA methylation, and the ability to refine structures using Monte Carlo minimization. The toolkit also provides geometric analysis tools based on rigid body formalism to evaluate DNA structures and trajectories. Together, these features enable users to model and analyze DNA configurations in high detail with a modular Python interface. By integrating structure generation and analysis into a single workflow, `MDNA` improves the study of DNA-protein interactions, paving the way for new insights into DNA dynamics and molecular simulations.

## 1 Introduction

Techniques such as X-ray crystallography, NMR, and cryo-EM are indispensable for obtaining high-resolution structures of biomolecules; however, they face limitations when applied to larger complexes or highly dynamic systems, where obtaining complete atomic detail and capturing transient states can be challenging. Predictive tools such as AlphaFold and RoseTTAFold have expanded our ability to model protein structures from sequence data Jumper et al. 2021; Abramson et al. 2024; Krishna et al. 2024, but reliable predictions are generally restricted to single proteins or smaller complexes, as accuracy depends heavily on the availability of high-quality input data, a significant limitation for large, heterogeneous assemblies. Molecular dynamics (MD) simulations provide a complementary approach, capturing the structure and dynamic behavior of biomolecules on biologically relevant timescales Hospital, J. R. Goñi, et al. 2015; Mohr et al. 2024. The setup of MD simulations requires tools that can generate molecular configurations and topologies in a consistent and flexible manner. Although structures for single proteins or small DNA-protein complexes can be derived experimentally or computationally, constructing larger molecular models containing DNA remains a significant challenge. Models of linear double-stranded DNA (dsDNA) of any sequence can be easily generated, but these approaches are inadequate for more intricate systems, such as protein-DNA complexes, DNA loops, or non-linear DNA topologies. Existing tools often lack the flexibility to construct such structures or to incorporate biologically relevant features like non-canonical bases, methylation patterns, or specific base pairing configurations.

Programs like Avogadro Hanwell et al. 2012 and UCSF ChimeraX Meng et al. 2023 allow users to build linear nucleic acid sequences. The user can set the number of base pairs per helical turn or instruct the software to generate the A- or B-form of dsDNA. PyMOL Schrödinger, LLC 2015 extends these functionalities by enabling chemical customization of DNA or RNA structures and integrating nucleic acids into existing molecular assemblies. The module DSSR (Dissecting the Spatial Structure of RNA) Lu 2020 is integrated into PyMOL, extending its capabilities for molecular visualization and structural analysis Zheng, Lu, and Wilma K Olson 2009; Li, Wilma K Olson, and Lu 2019. The proto-Nucleic Acid Builder (pNAB) supports the construction and geometry optimization of diverse nucleic acid analogs with alternative backbones and nucleobases Alenaizan et al. 2021. It is well-suited for generating initial models for MD simulations, particularly for small structures or novel backbone chemistries, though with limited spatial control. Although programs like Avogadro, UCSF ChimeraX, and PyMOL are well-suited for basic nucleic acid structure modeling, including base pair mutations and sequence-to-structure modules to generate linear DNA, they may be less adaptable for constructing more intricate DNA configurations with specific shapes or topological properties.

Building on the foundation of rigid body formalism to describe DNA conformations, F. Lankaš, Gonzalez, et al. 2009 another class of tools extends these principles to the construction and modeling of larger free-form DNA configurations. By treating each DNA base as a rigid entity with defined spatial positioning and orientation, these tools use geometric parameters to not only describe but also construct DNA structures. For example, the 3DNA software Lu and Wilma K Olson 2003, 2008 allows users to precisely reconstruct molecular structures using the complete set of rigid base parameters as input but lacks the features to specify shapes directly through these descriptors. Using mathematical descriptions of helical parameters, the VMD-based tool VDNA Bishop 2009 supports the creation of a range of structures from linear to circular DNA and even complex models such as nucleosome superhelices. However, the functionality of VDNA is confined primarily to a graphical user interface within VMD Humphrey, Dalke, and Schulten 1996, which can hinder reproducibility in large-scale studies. Alternatively, emDNA Young, Clauvelin, and Wilma K Olson 2022 is designed to model and optimize DNA loops and minicircles at the base pair level, integrating experimental configurations and considering sequence-dependent features to minimize elastic energy. Although emDNA provides tools to minimize energy and build DNA models at base pair resolution, it depends on 3DNA or DSSR to map optimized structures in atomic detail Lu 2020; Li, Wilma K Olson, and Lu 2019; Lu and Wilma K Olson 2003, which can complicate integration into broader simulation workflows.

On the other hand, coarse-grained models like oxDNA Ouldridge, Louis, and Doye 2011; Šulc et al. 2012; Rovigatti et al. 2015; Snodin et al. 2015; Poppleton et al. 2023 and tools like Polyply Grünewald et al. 2022 and `mrdna` Maffeo and Aksimentiev 2020 are designed to simulate DNA nanostructures and offer powerful capabilities for modeling large-scale systems, such as MD simulations of the entire minimal cell Grünewald et al. 2022; Stevens et al. 2023. In addition, OxViewBohlin et al. 2022, a tool for visualization and analysis of DNA nanostructures, provides an intuitive free-form editing tool to prepare models for simulation in the oxDNA engine, which users can export to other formats using tacoxDNA Suma et al. 2019 These tools, however, do not offer a solid analysis toolbox for detailed structural analysis of DNA conformations at all-atom resolution.

Instead, 3DNA Lu and Wilma K Olson 2003, 2008 and Curves+Lavery et al. 2009 have been the foundation for making the rigid body formalism accessible and consistent for analyzing nucleic acid structures and have significantly advanced understanding of DNA dynamics and flexibility F. Lankaš, Gonzalez, et al. 2009; Dršata and Filip Lankaš 2013. Subsequently, tools such as do_x3dna Kumar and Grubmüller 2015 and NAFlex Hospital, Faustino, et al. 2013 extend the capabilities of 3DNA and Curves+ by facilitating the analysis of MD trajectories and enabling calculations of DNA’s global helical axis, bending fluctuations, and elastic properties. Biobb_dna, part of the BioExcel building blocks ecosystem Andrio et al. 2019, aims to improve accessibility by providing parsing and processing for output files from tools such as Canal or Curves+ Lavery et al. 2009; Pasi, Maddocks, and Lavery 2015; Pasi, Zakrzewska, et al. 2017. SerraNA Noy and Golestanian 2012a; Velasco-Berrelleza et al. 2020 utilizes a Length-Dependent Elastic Model to analyze simulation data, focusing on DNA bendability and linking molecular fluctuations to macroscopic elastic behavior. Although all of these tools excel at analyzing nucleic acid structures and offer valuable insights into DNA dynamics, they are not designed primarily to generate large, heterogeneous assemblies containing DNA. In terms of analysis of DNA structure within MD simulations, libraries such as MDAnalysis Gowers et al. 2016, PyTraj Roe and Cheatham III 2013, and MDTraj McGibbon et al. 2015 provide general-purpose tools for trajectory analysis. Although they offer flexibility and efficiency in handling simulation data, they lack specialized functions for nucleic acid rigid body analysis. Although MDAnalysis previously contained a wrapper for 3DNA, this module is no longer maintained due to licensing restrictions Gowers et al. 2016. Other libraries like Bio3D Grant, Skjærven, and Yao 2021, HTMD Doerr et al. 2016, and VIAMD Skanberg et al. 2023 offer interactive and batch analysis capabilities, but do not specifically address the needs of nucleic acid simulations.

To address these challenges, we present `MDNA`: a software module for DNA structure generation and analysis. `MDNA` is a Python-based toolkit that integrates DNA structure generation and analysis into a cohesive framework suitable for molecular dynamics workflows. Our tool employs a spline-based mapping technique to generate accurate DNA structures using Monte Carlo (MC) minimization, supporting features such as the incorporation of non-canonical bases, manipulation of base pairing configurations (e.g., Watson-Crick-Franklin to Hoogsteen), addition of methylation patterns, and the extension or merging of existing DNA models. Furthermore, `MDNA` integrates post-simulation analysis tools based on a well-established rigid body formalism F. Lankaš, Gonzalez, et al. 2009, facilitating direct computation of DNA geometric properties within the MD simulation framework.

By consolidating structure generation and analysis tools into a single ecosystem, `MDNA` streamlines the process of preparing and analyzing DNA structures for molecular dynamics simulations. This integration facilitates the modeling of complex DNA systems that were previously challenging to construct and analyze, thereby expanding the capabilities of MD simulations in nucleic acid research. In the following sections, we detail the functionalities, applications, and advantages of the `MDNA` toolkit.

## 2 Results

### 2.1 Overview, Usage and Initialization of MDNA

The `MDNA` toolkit streamlines the construction of structural models of dsDNA and the analysis of dsDNA structures and molecular dynamics (MD) trajectories using the rigid base formalism F. Lankaš, Gonzalez, et al. 2009. `MDNA` serves a broad purpose, facilitating tasks from the calculation of rigid base parameters to the generation, editing, and building of complex DNA structures; see figure 1-a for a schematic data diagram containing an overview of methods. `MDNA` emphasizes flexibility, efficiency, and interoperability. To achieve these goals, `MDNA` leverages MDTraj to bridge MD data with the statistical analysis and scientific visualization ecosystem in Python McGibbon et al. 2015. MDTraj supports a wide range of MD data formats and calculations, providing a well-tested framework upon which `MDNA` builds. At the heart of `MDNA` is the `Nucleic` class, which unifies both the generation and editing of DNA structures and trajectory analysis. The `Nucleic` class contains the properties of the DNA structure along with its reference frames and trajectory information, providing methods for further manipulation and analysis.

**Figure 1:**
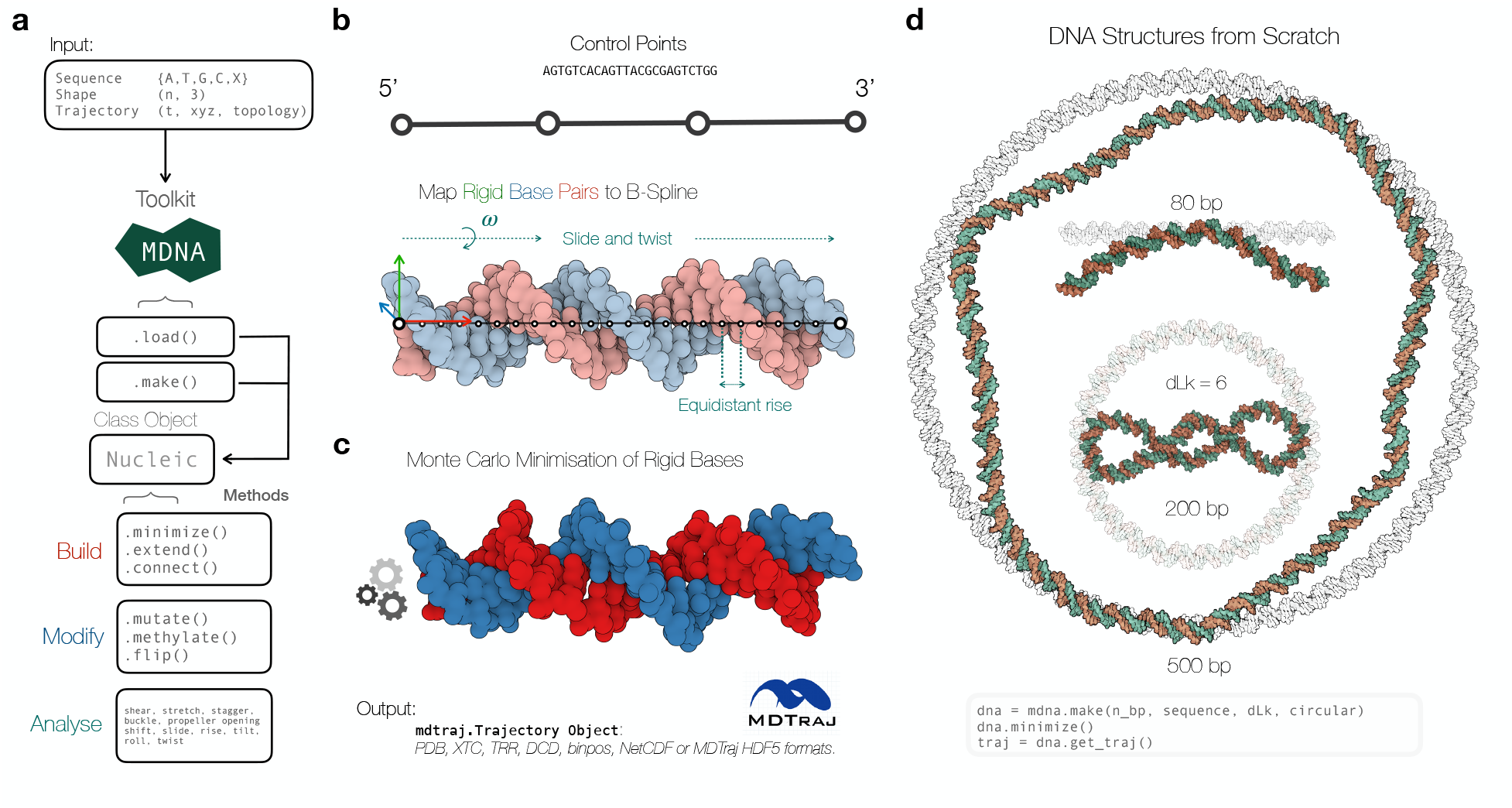
**a)** A workflow diagram depicting the capabilities of the `MDNA` toolkit. The diagram begins with input options—sequence (e.g., {A, T, G, C, X}, with X referring to non-canonical base codes.), shape, and trajectory—and proceeds to show the toolkit’s central class, `Nucleic`, which incorporates the functions `load()` and `make()` to initialize a `Nucleic` object. The diagram outlines how this object can be manipulated through categorized methods: *Build* (e.g., `minimize(), extend(), connect()`), *Modify* (e.g., `mutate(), methylate(), flip()`), and *Analyze* (e.g., measuring rigid base pair parameters). **b)** An illustration of mapping a sequence to a B-spline using control points, demonstrating how rigid base pairs are aligned to the spline to create an initial DNA structure. **c)** Visualization of the minimized DNA structure resulting from Monte Carlo minimization of rigid base steps, with output formats available via MDTraj-compatible formats (e.g., *pdb, xtc, trr*). **d)** Three examples of pre-minimized spline-based structures (transparent) compared with minimized structures (shown in orange and green): a 500 bp minicircle, an 80 bp linear strand, and a 200 bp minicircle with a `dLk` = 6, emphasizing writhe conservation. Molecular representations are visualized using Mol* Viewer Sehnal et al. 2021.

Initialization of the `Nucleic` class is achieved through the `mdna.make()` method, which generates a new DNA structure, or the `mdna.load()` method, which loads existing DNA structures or trajectories. Once initialized, the `Nucleic` class offers various methods such as `.minimize(), .mutate(), .methylate(), .flip(), .extend(), .connect()`. These methods allow users to manipulate DNA structures, introduce mutations, or join DNA fragments. The class also enables the retrieval of topological connectivity and rigid base parameters for structural analysis. For integration with other simulation tools, the `.get_traj()` method retrieves an MDTraj object of the current DNA configuration. This output can be used directly to start MD simulations in OpenMM Eastman et al. 2017, or can be exported to various formats, including *pdb, xtc, trr, dcd, binpos, netcdf, mdcrd, prmtop*, and more. In the following, we demonstrate how `MDNA` facilitates seamless integration of these functions into a cohesive workflow.

The `mdna.load()` function is the initial step for loading DNA structures and trajectories using MDTraj McGibbon et al. 2015 or base step reference frames along with a nucleotide sequence. For trajectory inputs, the function extracts the DNA sequence, determines the number of base pairs, and if the MDTraj trajectory contains multiple DNA strands, allows the user to specify the chain IDs of the strand of interest. Additionally, the circularity of the DNA structure is determined on the basis of the distance between 5’ and 3’ base pairs and a minimum sequence length, unless explicitly specified. For reference frame inputs, the function verifies the dimensionality and consistency of the data with the provided sequence, checking the number of time frames (`t`), the number of base pairs (`n_bp`), and the orthogonal triad (`t, n_bp, 4`). This initialization process ensures a consistent DNA representation and returns a `Nucleic` object.

### 2.2 Structure Generation and Minimization

DNA structure generation in `MDNA` is guided by a flexible spline-mapping protocol (see the Methods section 5.1 and 5.2) that enables the creation of atomic-resolution models of varying complexity. Using geometric input such as control points, sequences, and topological properties, `MDNA` provides users with precise control over DNA shapes, ranging from linear strands to complex loops. Energy minimization is integrated as an optional feature, ensuring biologically relevant configurations through Monte Carlo optimization (see the Methods section 5.3). This capability opens avenues for modeling unique and non-trivial DNA structures for molecular dynamics simulations.

The core function to initialize the `Nucleic` class is `mdna.make()`, which can generate DNA structures from scratch. This method allows flexibility and precision in the production of DNA models by accepting optional input arguments for nucleotide `sequence`, the number of base pairs (`n_bp`), topology (`circular`), adjustment in linking number (`dLk`), and shape definition (`control_points`). The `control_points` are sets of xyz coordinates used to determine the shape of a spline curve by specifying a series of interpolation points through which the curve passes. Input can be a custom set of points from a parametric function that describes the desired shape (e.g., an ellipse, superhelix, trefoil knot, etc.), with the only requirement that at least four control points for the interpolation of the spline curve be provided. Once the spline is fitted through the control points, `mdna.make()` equidistantly distributes the positions of the base pairs along the spline using a spacing of 0.34 nm, which corresponds to the average rise between base pairs. The base pair positions and the tangents of the spline’s space curve are used to construct the base pair reference frames (see the Methods section 5.1). Starting from the 5’ end, the subsequent base pair reference frames are twisted with respect to the previous base pair to ensure each turn of the double helix contains 10.5 base pairs, resulting in the Darboux frame representation of the DNA structure (see the Methods section 5.2). The user can modify the number of turns per base pair. Circular DNA has a closed topology and may have excess or reduced twist, which is distributed over all base pairs in the sequence. Using the input sequence, the atomic reference coordinates are assigned to the Darboux frame, generating the topological structural information to initialize an MDTraj object (see the Methods section 5.2). All input arguments are optional; if only a DNA sequence is provided, the default settings generate a linear DNA geometry. Users can also control the number of base pairs to scale the shape and adjust the twist via the linking number difference (`dLk`), see the Methods section 5.5 for more details on linking numbers in DNA. This functionality ensures that the generated DNA structures can be tailored to specific experimental or theoretical requirements. If a closed topology is desired without a specified shape, a circular configuration is generated.

The `Nucleic` instance from `mdna.make()` contains by default B-DNA geometries without thermal fluctuations as specified by the control points. To optimize and minimize the energy of the given structure, the `nucleic` instance contains the `.minimize()` function, which is designed to relax the DNA structure using Monte Carlo (MC) simulations (see the Methods section 5.3 for implementation details). This relaxation process targets a low-energy configuration by minimizing elastic deformations and steric clashes, modeled through an elastic Hamiltonian and hard sphere potentials. Users can tune the minimization process by adjusting the temperature, excluded volume parameters, and fixing specific base orientations. For example, by keeping the position and orientation of the ends of the DNA strand fixed or by providing a list of base step indices whose orientation and position remain fixed during minimization. When importing a trajectory, a selected snapshot (or time point) can be minimized using the MC model. By default, we aim for convergence of the system using an exponential decay model for the energy, to avoid being trapped in a local minimum. If topological properties such as the linking number need to be conserved, the energy is minimized. Equilibrating the writhe relies on checking whether the energy starts to fluctuate around a fixed value, as this process involves a barrier crossing. Figure 1-b shows a visual illustration of the spline mapping to construct a DNA structure, as well as the second (optional) stage of MC energy minimization (figure 1-c). Finally, Figure 1-d presents example DNA structures generated using `mdna.make()` and subsequently minimized with the MC module. The examples include an 80 bp linear strand, a 200 bp minicircle with `dLk` set to 6, and a 500 bp minicircle. In the minimized structure of the 200 bp minicircle, we clearly observe the crossings (writhe) in the DNA, which is the result of the conservation of the change in linking number during the MC minimization. The computational cost of structure generation and minimization up to 1000 base pairs is negligible and can be done within a minute on an everyday notebook.

### 2.3 Sequence Library and Nucleotide modifications

`MDNA` provides an extensive sequence library that supports the modeling of canonical and non-canonical nucleobases, enabling the exploration of diverse structural properties of DNA. This feature allows researchers to introduce mutations, methylation patterns, and novel base analogs into their models, facilitating studies on topics ranging from genetic mutations to synthetic biology. The toolkit’s library extends to fluorescent and hydrophobic bases, as well as orthogonal systems like Hachimoji DNA. The `.mutate()` module changes nucleobases in a DNA structure, supporting both complementary and non-complementary pairings. The module includes the canonical bases adenine (A), thymine (T), guanine (G), cytosine (C), and uracil (U) as well as various noncanonical base analogs. The Hachimoji DNA system is available, consisting of four extra synthetic nucleotides forming orthogonal pairs: B-S and P-Z (PDB: 6MIG, 6MIK, 6MIH) Hoshika et al. 2019. The library also includes the hydrophobic base pair d5SICS - dNaM Malyshev et al. 2014, an example of unnatural base pairs that interact without hydrogen bonds (PDB: 3SV3, 4C8M) Betz, Malyshev, Lavergne, Welte, Diederichs, Dwyer, et al. 2012; Betz, Malyshev, Lavergne, Welte, Diederichs, Romesberg, et al. 2013. Furthermore, two fluorescent bases: 2-aminopurine Ward, Reich, and Stryer 1969 and the tricyclic cytosine analog are available in the nucleotide sequence library Wilhelmsson et al. 2003, both noted for their fluorescent properties in different environments essential in fluorescence studies in nucleic acid research Xu, Chan, and Kool 2017 (PDB: 2KV0, 1TUQ) Dallmann et al. 2010; Engman et al. 2004. Note that parameters for all xenogenic nucleobases are not yet available in MD force fields, as this is an ongoing community effort. However, general force fields, such as GAFF(2) and CGenFF He et al. 2020; Vanommeslaeghe and MacKerell Jr 2012, are available and can often be sufficient. Alternatively, specific parameters can be derived for individual nucleobases; for example, the parameters of 2-aminopurine (2AP) are available and are based on quantum chemical DFT calculations Remington, McCullagh, and Kohler 2019. See Table 1 for an overview of all available nucleobases in `MDNA`.

**Table 1:** List of available nucleobases with type, name, code and complementary base pairing matches.

| Name | Code | Complementary | Type |
| --- | --- | --- | --- |
| Adenosine | A | T | Canonical |
| Thymine | T | A | Canonical |
| Guanine | G | C | Canonical |
| Cytosine | C | G | Canonical |
| Uracil | U | A | Canonical |
| 2-aminopurine | E | T | Fluorescent |
| Tricyclic cytosine | D | G | Fluorescent |
| A-analogue | B | T,D | Hachimoji |
| T-analogue | D | A,B | Hachimoji |
| G-analogue | Z | C,P | Hachimoji |
| C-analogue | P | G,Z | Hachimoji |
| d5SICS | L | M | Hydrophobic |
| dNaM | M | L | Hydrophobic |

The `.flip()` command rotates the nucleobases around the glycosidic bond that connects the nucleobase to the DNA sugar backbone. By default, nucleobases rotate 180 degrees, shifting from a Watson-Crick-Franklin (WCF) to a Hoogsteen (HG) configuration. The HG base pairing motif, an increasingly recognized structural property of DNA structures, involves a rotation of purine from an anti-to syn orientation Hoogsteen 1959. In this geometry, 5-ring purine, rather than the 6-ring, forms a hydrogen bond with the pyrimidine. HG configurations are more common in AT base pairs, with GC pairs requiring protonated cytosine to accommodate this bonding pattern Nikolova et al. 2013. The function `.methylate()`allows users to methylate C or G bases, with the option to automatically methylate cytosines at the CpG sites, where most DNA methylation occurs Moore, Le, and Fan 2013. Figure 2-a shows the extensive sequence library in `MDNA`, featuring the canonical dsDNA bases, nucleotide modifications and artificial bases. The molecular representations visualized highlight the toolkit’s ability to handle a broad range of DNA structural variations, aiding researchers in exploring the effects of these structural modifications. Both the `.flip()` and `.methylate()` only require a list of nucleobase indices as input to indicate which base to flip or methylate.

**Figure 2:**
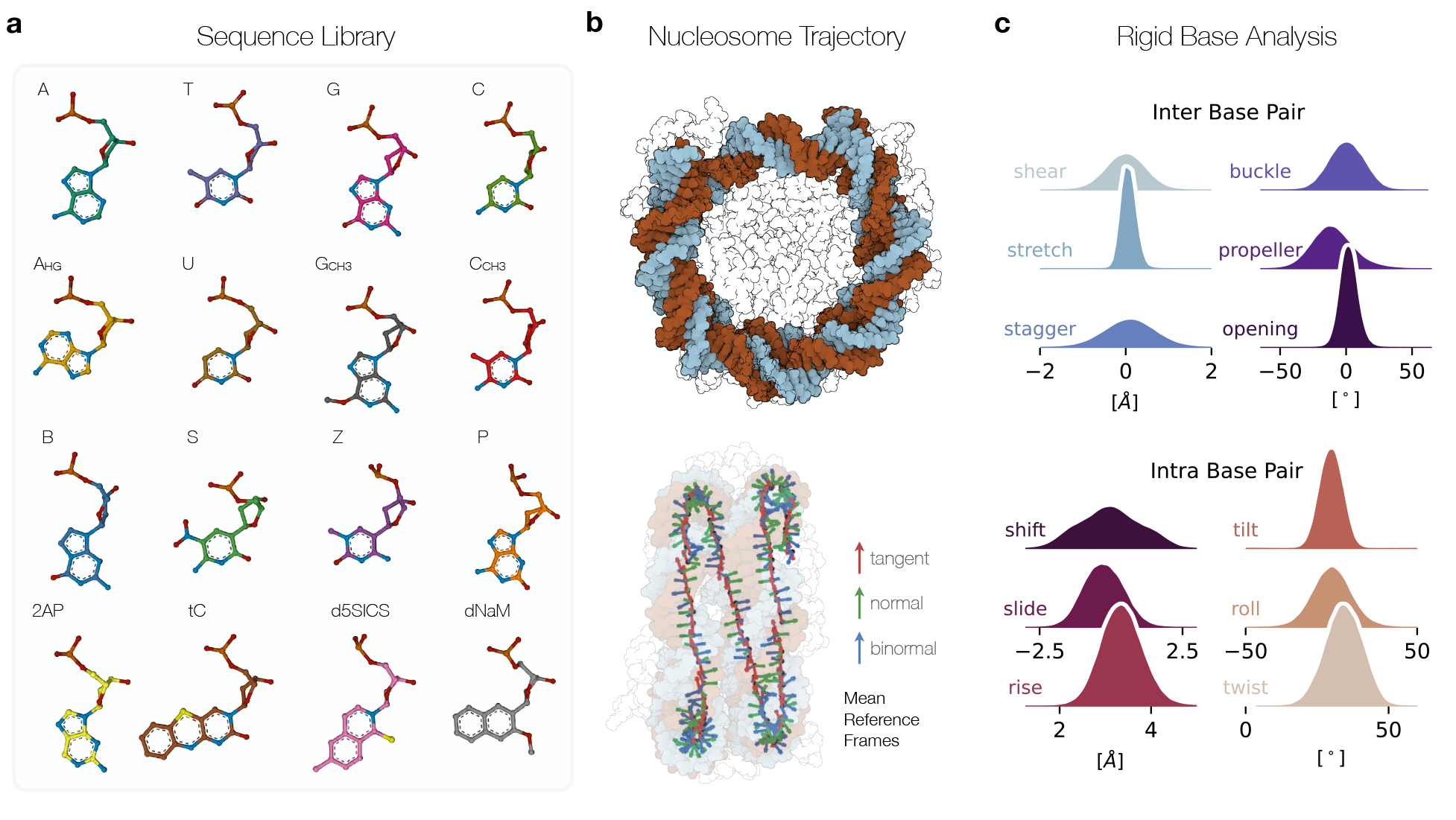
**a)** Sequence library showing molecular representations of the available nucleobases mutations and modifications in the `MDNA` toolkit. Each nucleobase is annotated with its letter code or acronym. The first row shows the 4 canonical dsDNA bases, the second row Adenosine in Hoogsteen conformation, followed by Uracil, methylated Guanine, and Cytosine. The third row shows the Hachimoji base analogs: B, S, Z, and P respectively. The last row shows two fluorescent nucleobases 2-Amino Purine (2AP) and tri cytocine (tC), and the hydrophobic nucleobase set d5SICS and dNaM. **b)** Front view of a nucleosome core particle with 147 bp DNA and side view of the Darboux frame of the DNA wrapped around the histone showing the mean reference frame of each basepair (PDB 1KX5 Davey et al. 2002). **c)** Cumulative rigid base parameter distributions computed with *MDNA* of the nucleosome trajectory of 100 ns. Molecular representations are visualized with Mol* Viewer Sehnal et al. 2021.

### 2.4 Rigid Base Analysis Modules

We offer the first native Python implementation to calculate the rigid base parameters of the structural structure. For intra-base pair movements, they detail rotations (buckle, propeller, opening) and translations (shear, stretch, stagger), capturing the nuances of individual base pair positioning. For inter-base pair steps, they define rotations (tilt, roll, twist) and translations (shift, slide, rise), providing the spatial and rotational relationships between adjacent base pairs, also known as step parameters. Previously, this required software installations compiling with x3DNA, Curves+/Canal or the dependency on web services for MD trajectory analysis Kumar and Grubmüller 2015; Pasi, Zakrzewska, et al. 2017. Users can extract these parameters by providing an MDTraj object in the `mdna.compute_rigid_parameters()` function or use `Nucleic` class *getters* to obtain a Numpy array (shape: `t, n_bp`, 12) representing the relative translation and rotation between each base pair, with `t` the number of trajectory time frames, and `n_bp` the number of base pairs. The Python implementation uses vectorized arrays using numpy Harris et al. 2020. We used the theoretical framework described in Curves+ Lavery et al. 2009 and Petkevičiūtė et al. to calculate the rigid base parameters Petkeviciute 2012. See the Methods section 5.4 for the calculation of rigid parameters. Figure 2-b and c illustrate the analysis of rigid base parameters of an MD trajectory of 250 ns of the Nucleosome Core Particle (PDB 1KX5). Rigid base parameters enable the calculation of global DNA properties, such as the persistence length, which indicates the stiffness of the DNA derived from correlations in base pair orientations; higher correlations suggest a greater stiffness of the DNA strand Mazur 2007; Enrico Skoruppa, Laleman, et al. 2017. Furthermore, global tilt and roll parameters help to compute the total curvature of the DNA helix, providing information on the flexibility of the DNA Strahs and Tamar Schlick 2000. In addition to the rigid base parameters, `MDNA` includes the `mdna.compute_linking_number()` method to calculate the linking number, an important topological property of DNA (see the Methods section 5.5). All of these properties are essential structural metrics for understanding the supercoiling, bending, and interactions of DNA with proteins.

### 2.5 Building Biomolecular Assemblies

The ability to construct DNA-protein assemblies highlights `MDNA`’s strength in enabling advanced modeling scenarios. Case studies include integration of DNA strands with H-NS filaments, extension of nucleosome-bound DNA to linker regions, and the creation of protein-mediated DNA loops. These examples demonstrate the utility of the toolkit in the construction of biologically relevant systems and bridge the gap between structural data and computational modeling studies. In the first example (Figure 3-a), the `.make()` function is used to create a continuous DNA strand along an H-NS filament. Starting with a filament of 12 H-NS subunits Van Der Valk et al. 2015, four DNA Binding Domains (DBDs) are selected, and a 12 bp DNA strand is superimposed onto these domains Riccardi et al. 2019; Heesch, Bolhuis, and Vreede 2023. See the Methods section 5.6 for details on how the H-NS filament was constructed. This method allows for the generation of a new DNA strand that spans the entire length of the filament, showcasing the tool’s ability to construct specified DNA shapes from scratch using protein filaments as scaffolds.

**Figure 3:**
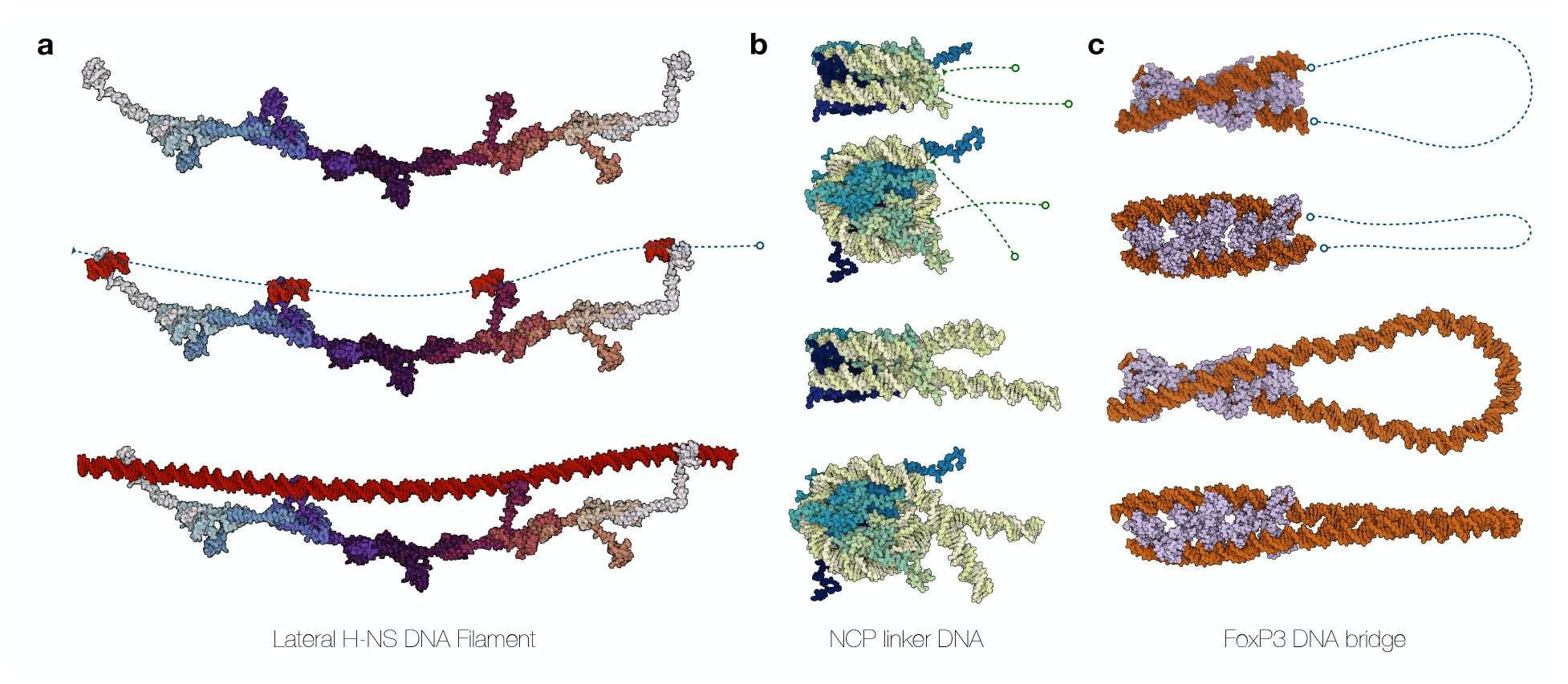
**a**) Constructing a continuous DNA strand along the H-NS filament. The process begins with an H-NS filament consisting of 12 subunits Van Der Valk et al. 2015; Valk et al. 2017. Next, four DNA Binding Domains (DBDs) positioned along one side of the filament are selected, and a 12 bp DNA strand is superimposed onto these domains using structural data from the H-NS-DNA complex (PDB: 1HNS Shindo et al. 1995) Riccardi et al. 2019; Heesch, Bolhuis, and Vreede 2023. The DNA oligomers are shown in red. Finally, the reference frames of the four DNA oligomers are used as control points to generate a new DNA strand that spans the entire length of the H-NS filament. **b**) Nucleosome (PDB 1KX5 Davey et al. 2002) extended with 36 bp linker DNA in both the 5’ and 3’ direction. The extended DNA sequences are energy-minimized using the MC module. **c**) Higher-order FoxP3 multimer that forms a ladder-like bridge to DNA (PDB 8SRP)Zhang et al. 2023. Here we show how the bridged DNA ends can be connected with 200 bp that are minimized using the `.minimize()` function, creating a DNA loop held together by the FoxP3 ladder. Molecular representations are visualized with Mol* Viewer Sehnal et al. 2021.

In the second example (Figure3-b), the `.extend()` function is employed to extend a DNA sequence, adding 36 bp linker DNA to a nucleosome in both the 5’ and 3’ directions. This function accepts several parameters that allow precise control over the extension of the DNA strand: Users can specify the number of base pairs (`n_bp`) or provide a DNA sequence for the extension. The `forward` boolean parameter determines the direction of the extension. Users can also choose a time frame (`frame`) and shape (`control_points`) to define the new DNA shape. Users can determine if the terminal base pairs of the original structure are included or excluded from the minimization procedure (`margin`); increasing this parameter includes more bases of the original strand to also be minimized. This feature enables smoother transitions between the original DNA and the newly generated DNA.

The `.connect()` function is able to create a single DNA structure from two separate segments, and similarly to `.extend()` includes the arguments `n_bp`, `frame, control_points`, and `margin`. The function interpolates a straight line between the ends and distributes the optimal number of base pairs to achieve a neutral twist. The function requires two `Nucleic` objects and optional parameters to generate the DNA structure. In the final example (Figure 3-c), the `.connect()` function is used to link two separate DNA structures by creating a strand between the 3’end of the first and the 5’end of the second. This example illustrates how the function can generate a continuous DNA loop held together by the FoxP3 protein, which forms a ladder-like bridge between the two DNA ends Ramirez et al. 2022; Z. Liu et al. 2023. Together, the functions `.extend()` and `.connect()` further provide additional utility, allowing the creation and manipulation of complex DNA structures with precision and ease. These capabilities advance the study of molecular simulations in large, complex heterogeneous systems.

## Discussion

`MDNA` advances molecular simulations by providing atomic-resolution structural modeling of double-stranded DNA (dsDNA) in diverse shapes and compositions, including DNA-protein assemblies. By allowing for precise structural modeling of DNA at atomic resolution, `MDNA` contributes to improving our understanding of DNA dynamics and interactions in complex biological systems. The integration of structure generation, editing, and analysis tools into a single platform provides researchers with a streamlined workflow to address challenging questions in structural biology and molecular simulation. Its core strength lies in a spline mapping protocol that enables the construction of template DNA structures in arbitrary shapes, addressing limitations in existing tools that often require extensive manual adjustments or are constrained in scope. In contrast, `MDNA` facilitates the generation of diverse configurations with high accuracy supported by a Monte Carlo (MC) minimization, including those incorporating non-canonical bases, Watson-Crick-Franklin (WCF) to Hoogsteen (HG) transitions, and specific methylation patterns.

The case studies presented in this work, such as the construction of DNA strands along an H-NS protein filament and the extension of nucleosome DNA, highlight `MDNA`’s capability to handle complex biological systems critical for studying DNA dynamics and protein interactions. By allowing initial configurations of atomic resolution tailored to specific experimental, theoretical, and topological needs, `MDNA` fills a significant gap in the existing toolkit for molecular simulations. In addition to its structure-generation capabilities, `MDNA` integrates advanced analysis tools within a unified Python-based ecosystem Van Rossum and Drake Jr 1995. The integration of geometric analysis modules based on the rigid body formalism in the toolkit F. Lankaš, Gonzalez, et al. 2009 further improves the utility to understand the structural dynamics and interactions of DNA. By incorporating libraries such as MDTraj and OpenMM McGibbon et al. 2015; Eastman et al. 2017, `MDNA` consolidates structure generation, simulation, and analysis into a streamlined workflow, reducing dependence on disparate software platforms and improving accessibility for researchers at varying levels of expertise.

While `MDNA` excels in generating customizable structures and integrating analysis tools, some limitations present opportunities for future improvement. The lack of direct support for single-stranded DNA (ssDNA) structures restricts its applicability to processes such as replication, transcription, certain DNA repair pathways, and RNA modeling, which often involve hybrid or single-stranded configurations. Although ssDNA can be approximated by modifying the twist and removing the antistrand, a dedicated module for ssDNA and RNA generation would greatly enhance the versatility of the toolkit, particularly for studying hybrid RNA-DNA structures and RNA folding dynamics. The current MC-based structure relaxation, while effective, is the only optimization tool currently integrated into MDNA; other methods, such as steepest descent, would improve the flexibility of the toolkit. Additionally, the simulation of structures with the sequence library relies on existing force field parameters, which may not be available or adequately parameterized for non-canonical bases and modified nucleobases. These non-canonical bases are also incompatible with minimization using the MC model, underscoring the ongoing challenges in molecular modeling. `MDNA` is not yet optimized for kilobase (kb) length DNA structures; improving scalability for larger systems is a feasible and promising future direction. The optional nature of the minimization is advantageous for certain applications, as `MDNA` allows users to specify initially energetically unfavorable configurations, such as very tight knots or tiny minicircles. This flexibility cautions that initial structures should be used for qualitative analysis, as they may be nonphysical depending on the choice of control points and may deviate from physical realism without further refinement. Future updates could also explore the integration of enhanced sampling methods, such as OpenPathSampling Swenson et al. 2018 or Plumed Tribello et al. 2014 workflows, to optimize starting structures and analyze free-energy landscapes in high-throughput scenarios.

In conclusion, `MDNA` is a powerful and versatile tool for the generation of dsDNA structures that provides detailed insight into various dsDNA architectures and improves the setup of molecular dynamics simulations for DNA and protein-DNA complexes. The comprehensive features and seamless integration with existing MD tools make `MDNA` a useful resource for researchers. Future updates will focus on expanding the support for the sequence library, introducing ssDNA and RNA functionality, and improving optimization algorithms. With its comprehensive features and strong community focus, `MDNA` establishes itself as an essential tool to advance DNA structure modeling and analysis. To support its adoption, `MDNA` is complemented by extensive tutorials, demos, and a detailed GitHub repository and documentation page: https://github.com/Heezch/mdna. These resources improve accessibility for novice and experienced users, providing a foundation for educational applications such as workshops or classroom demonstrations.

## Acknowledgements

This publication is part the OCENW.KLEIN.200 grant (T.v.H. and J.V.) and financed by the Dutch Research Council (NWO). ES, and HS were supported by the Deutsche Forschungsgemeinschaft (DFG, German Research Foundation) under Germany’s Excellence Strategy - EXC-2068 - 390729961. The codebase (MIT License) is available at https://github.com/Heezch/mdna.

## 5 Methods

### 5.1 Frame Generation Algorithm

This section explains how to compute coordinate frames along a spline. A related problem in mathematics and computer graphics is the generation of coordinate frames with minimal torsion along a spline Bloomenthal 1990. The solution involves initializing a coordinate frame at the beginning of the curve and incrementally propagating this frame along the spline Bloomenthal 1990. This method relies only on the first derivative of the curve to compute subsequent frames and avoids problems with inflection points and torsion by maintaining continuity based on the previous frame. The process begins with the generation of a space curve, which serves as the helical axis of the DNA. The next step involves computing coordinate frames along equidistantly spaced points on the curve to guide the placement of the DNA’s base pairs. Finally, a twist can be introduced to obtain a helical structure.

Given a set of control points **P** = *P*_1_, *P*_2_, …, *P*_*n*_ in ℝ^3^, a spline curve *S* : [0, 1] → ℝ^3^ is defined as a piecewise polynomial function that smoothly interpolates through these points. The spline is parameterized by a non-dimensional parameter *u*, where *u* ∈ [0, 1], represents the normalized arc length along the spline. For a cubic B-spline, the spline curve *S*(*u*) as a weighted sum of control points *P*_*i*_ at any point *u* can be expressed as:

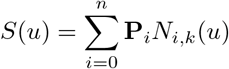

where *N*_*i,k*_(*u*) are the basis functions of the B-spline with degree *k*, and **P**_*i*_ are the control points De Boor 1978. The degree is set by default to *k* = 3.

The procedural generation of DNA structures involves computing coordinate frames along *S*(*u*), beginning with the computation of an initial frame followed by the propagation of this frame along the spline Bloomenthal 1990. The goal is to construct an orthonormal basis defined by T, N, and B, which establishes a positively oriented moving frame at each curve point.

The initial tangent vector *T*_0_ is calculated as the normalized first derivative of *S*(*u*) at the starting point, indicating the direction of the curve:

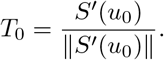

The initial normal vector *N*_0_ is determined by crossing *T*_0_ with a predefined *reference* vector **v**_*ref*_, typically initialized as [0, 0, 1]^*T*^, and normalizing this cross product. If *T*_0_ is parallel to **v**_*ref*_, an alternate vector is selected to ensure a non-zero cross product:

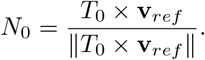

The initial binormal vector *B*_0_ is calculated as the cross product of *T*_0_ and *N*_0_:

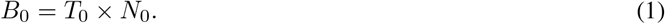

For each subsequent point *u*_*i*_ along the spline, the tangent vector *T*_*i*_ at *u*_*i*_ is computed as the normalized derivative of *S*(*u*):

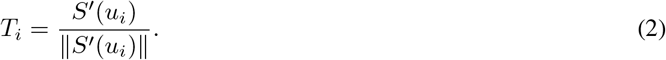

The axis of rotation **a**_*i−*1,*i*_ is determined as the normalized cross product of *T*_*i−*1_ and *T*_*i*_, indicating the direction to rotate *N*_*i−*1_ to compute *N*_*i*_ :

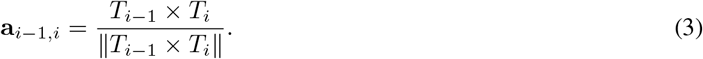

If the norm of **a**_*i*−1,*i*_ is very small (indicating negligible rotation), *N*_*i*_ is set to *N*_*i*_−_1_ to avoid discontinuity of the minimal torsion in the reference frames:

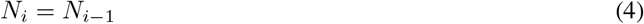

Otherwise, the angle of rotation *θ*_*i−*1,*i*_ is computed using the dot product of *T*_*i−*1_ and *T*_*i*_ :

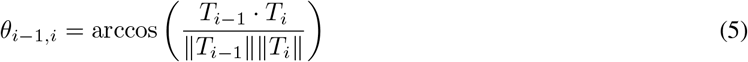

Using the computed axis **a**_*i*−1,*i*_ and angle *θ*_*i*_ −_1,*i*_, *N*_*i*_ is obtained by rotating *N*_*i*_− _1_ around **a**_*i*_− _1,*i*_. Finally, at *u*_*i*_ the binormal vector *B*_*i*_ is computed as:

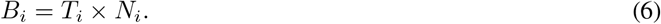

The result is a twist free geometry described by an orthonormal coordinate system at a point on a surface or curve, consisting of tangent, normal, and binormal vectors that describe local geometric properties, which is also known as a Darboux frame. A practical implementation of constructing and evaluating B-splines and their derivatives is provided by the `scipy` library Virtanen et al. 2020.

Note that the more conventional approach using Frenet-Serret frames encounters difficulties at inflection points, where the second derivative of the curve is zero Pressley 2010. At these points, the normal and binormal vectors become undefined, leading to discontinuities and sudden inversions in the orientation of the coordinate frames. Although some approaches attempt to address this by introducing a fixed reference vector within the global coordinate system, these can still result in unwanted twisting at inflection points or regions where the curve aligns with the reference vector.

The spline generation and frame computation are the basis of our method for the procedural generation of DNA structures. To create a Darboux frame that describes DNA as a space curve, we can introduce a helical twist by simply rotating the consecutive basis vectors along the base of **T**(*u*) by the number of base pairs per turn (typically 10.5 bp). Alternatively, we can locally adjust the twist by specifying a custom twist value for a defined range of base pairs. The parameter *u* is evenly distributed along the spline, with all segments having the same size, *d*, which by default corresponds to an increase value of 0.34 nm. The distribution of frames is determined by parameterizing the arc length of the spline. The goal is to distribute *n* points along the spline, with fixed positions at both ends and a default spacing of 0.34 nm between each point. Due to the end constraints, the spacing of the last segment may be slightly larger than or less than 0.34 nm. To mitigate this, the total difference across the complete sequence is evenly distributed.

### 5.2 Back Mapping of Base Pairs

The methodology for back mapping base pairs uses the DNA’s Darboux frame, explained in Section 5.1, as a template to construct the atomic coordinates. The back mapping of the frames to the atomic coordinates starts with determining the sequence from the generated frames. After the sequence is set, the MDTraj library is used for the initialization of the trajectory, which contains the xyz coordinates and the topology of the system McGibbon et al. 2015.

Initialization of the trajectory involves the generation of dummy coordinates for the sequence and the use of the Tsukuba convention Dickerson 1989 to add a reference base frame to each base, using ideal forms for the initial structure of the bases. Subsequently, for the antisense chain, complementary bases are identified for positions from *N* to 2*N−* 1, and their coordinates undergo a transformation to ensure proper base pairing. This transformation involves rotating and flipping the coordinates of each complementary base by 180 degrees around the x-axis, which lies in the plane of the base. This procedure guarantees that the first base of the anti-sense chain complements the sense strand’s first base. The topology construction adheres to the progression from 5’ to 3’ for the leading chain with the anti-sense chain in reverse orientation. For circular DNA structures, we ensure the connection of terminal ends in their respective chains.

Following the preliminary construction of the DNA trajectory and topology with dummy coordinates, we compute the mean reference frames for the base pairs as described in Section 5.4. This step allows for the subsequent update of coordinates based on the Darboux frame derived from a spline. Let **O**_dummy_ and **O**_new_ denote the origins of the dummy and the new frames, respectively, and **B**_dummy_ and **B**_new_ represent the bases of these frames. Transformation from the old frame to the new frame involves calculating the rotation **R** and translation **T** matrices. The rotation matrix **R** is obtained by solving **B**_new_ = **RB**_dummy_, and the translation is determined by **T** = **O**_*new*_ *−***O**_dummy_. These transformations are then applied to the dummy coordinates, finalizing the structural model with **X**_new_ = **RX**_dummy_ +**T**, where **X**_dummy_ and **X**_*new*_ denote the sets of old and updated coordinates, respectively.

### 5.3 Monte Carlo Minimization

∈

We provide energy minimization and thermalization of the generated DNA structures in the form of Monte Carlo (MC) sampling based on the rigid base pair framework F. Lankaš, Gonzalez, et al. 2009; Eslami-Mossallam et al. 2016. Using the procedure described in 5.1 and 5.4, DNA molecules are mapped onto a sequence of triads. Relative orientations and positions of neighboring triads are parametrized in terms of the 3 rotational degrees of freedom (tilt, roll, and twist) and 3 translational degrees of freedom (shift, slide, and rise) that are condensed into junction vectors **X**_*i*_ −ℝ^6^. These vectors are then decomposed into static and dynamic components

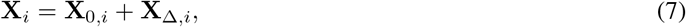

where the static components **X**_0,*i*_ specify the ground state of the molecule. The most important components of this vector are the intrinsic twist, reflecting the helicity of the molecule, and the intrinsic rise that determines the contour length per step of base pair. Fluctuations around this ground state are specified by an elastic Hamiltonian ℋ (**X**_Δ_) that penalizes deformations with the harmonic energy F. Lankaš, Gonzalez, et al. 2009; W. K. Olson et al. 1998; F. Lankaš, Šponer, et al. 2003; Enrico Skoruppa and Schiessel 2025

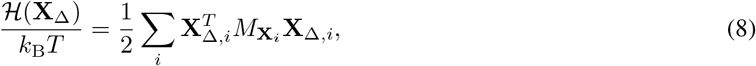

where 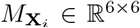 are sequence-dependent stiffness matrices and the summation extends to all junction vectors.

Finally, we denote *k*_B_ as the Boltzmann constant and *T* as the temperature. At present, stiffness matrices and ground states are taken from atomistic simulations F. Lankaš, Šponer, et al. 2003 or crystallographic data W. K. Olson et al. 1998. In this parametrization, the elastic energy is assumed to be local, i.e., fluctuations at each base pair step are decoupled from neighboring steps. Work of the last decade suggests this locality be insufficient to capture detailed length-scale dependent elastic features Noy and Golestanian 2012b; Enrico Skoruppa, Voorspoels, et al. 2021; Segers et al. 2022. Parameters derived from more recent models (e.g., cgNA+ Sharma et al. 2023) indeed provide a more general elastic description featuring non-local couplings. To thermalize and minimize DNA structures as initial configurations for Molecular Dynamics simulations, the local description is deemed sufficient.

Steric hindrance and screened electrostatic repulsion are modeled using hard-sphere potentials with effective diameters *d*_*EV*_, depending on the ionic conditions of the experiment Rybenkov, Cozzarelli, and Alexander V Vologodskii 1993. At a concentration of 150 mM univalent salt, an effective diameter of *d*_*EV*_ = 4.0 nm was found to agree well with the experimental single-molecule measurements Vanderlinden et al. 2022. Configurations are progressively generated with a series of permutation moves. These include pivot, crankshaft, and single-triad permutation (rotation and translation) moves Vanderlinden et al. 2022; E. Skoruppa and Carlon 2022, see Figure 4 for an illustration of these MC moves. The type of move and the number of triads involved are chosen at random. Rejection and acceptance are based on the Metropolis criterion.

**Figure 4:**
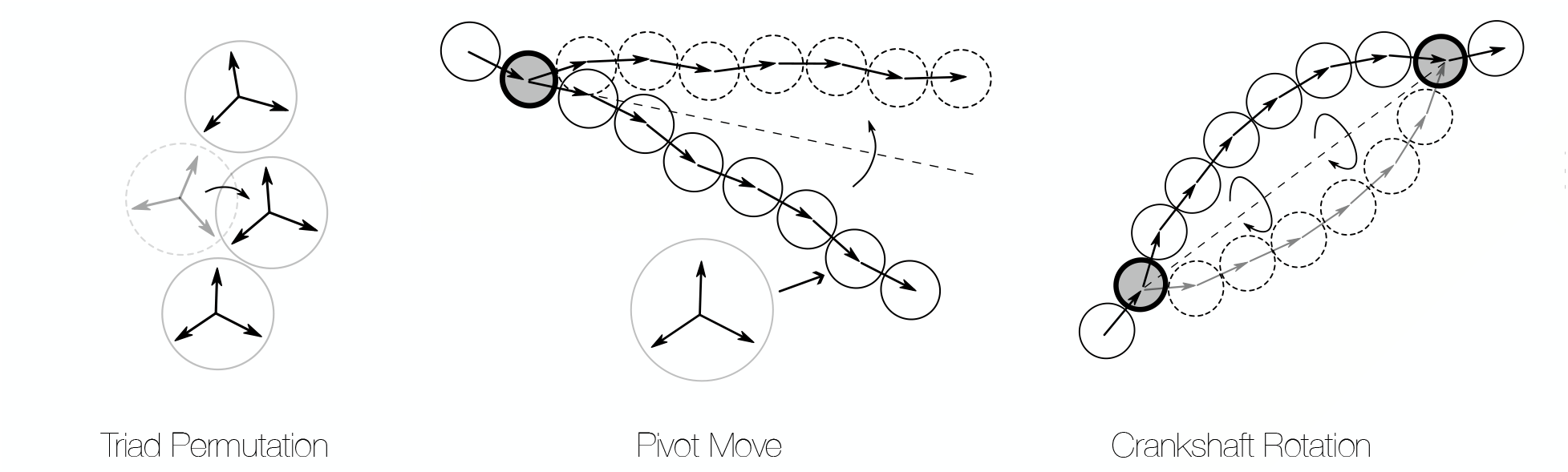
The representation of moves used in the DNA triad model for generating new configurations using the MC method is illustrated from left to right as follows: single triad permutation, pivot move, and crankshaft rotation.

### 5.4 Calculation of Rigid Parameters

The Cambridge convention established the rigid body parameters for the geometry of nucleic acids but does not specify the precise construction of the reference frame Dickerson 1989. Tools such as 3DNA Lu and Wilma K Olson 2003 and Curves+ Lavery et al. 2009 demonstrate the effectiveness of rigid body models, serving as benchmarks for the rigid base definitions, with their similarities and differences extensively described elsewhere Lankas and Schlick 2012. Rigid body models treat each DNA base as a single entity. Each base *b* is assigned a reference frame **B**_*b*_ for precise spatial positioning and orientation. The frame consists of a reference point and an orthonormal triad of vectors pointing towards the major groove, backbones, and directing perpendicularly to the base plane. The collection of these triads is often called the Darboux frame. Together transformations of the Darboux frame can distinguish DNA conformations using rotational matrices and translation vectors for both intra- and inter-base pair coordinates. For intra-base pair movements, they detail rotations (buckle, propeller, opening) and translations (shear, stretch, stagger), capturing the nuances of individual base pair positioning. On the other hand, for inter-base pair steps, they define rotations (tilt, roll, twist) and translations (shift, slide, rise), providing the spatial and rotational relationships between adjacent base pairs, also known as step parameters.

To obtain the rigid base parameters we followed the implementation as described in Curves+ Lavery et al. 2009 and Petkevičiū tė et al. Petkeviciute 2012. For each base, we construct a reference frame 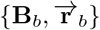 consisting of an orthonormal triad of unit vectors **B**_*b*_ = {*ê*_0_, *ê*_1_, *ê*_2_} and their origin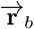. In this setup, the base triad vector *ê*_0_ points to the major groove, *ê*_1_ connects the backbones, and *ê*_2_ is normal to the base plane. See the top panel of Figure 5-a for a cartoon representation of a single base triad.

**Figure 5:**
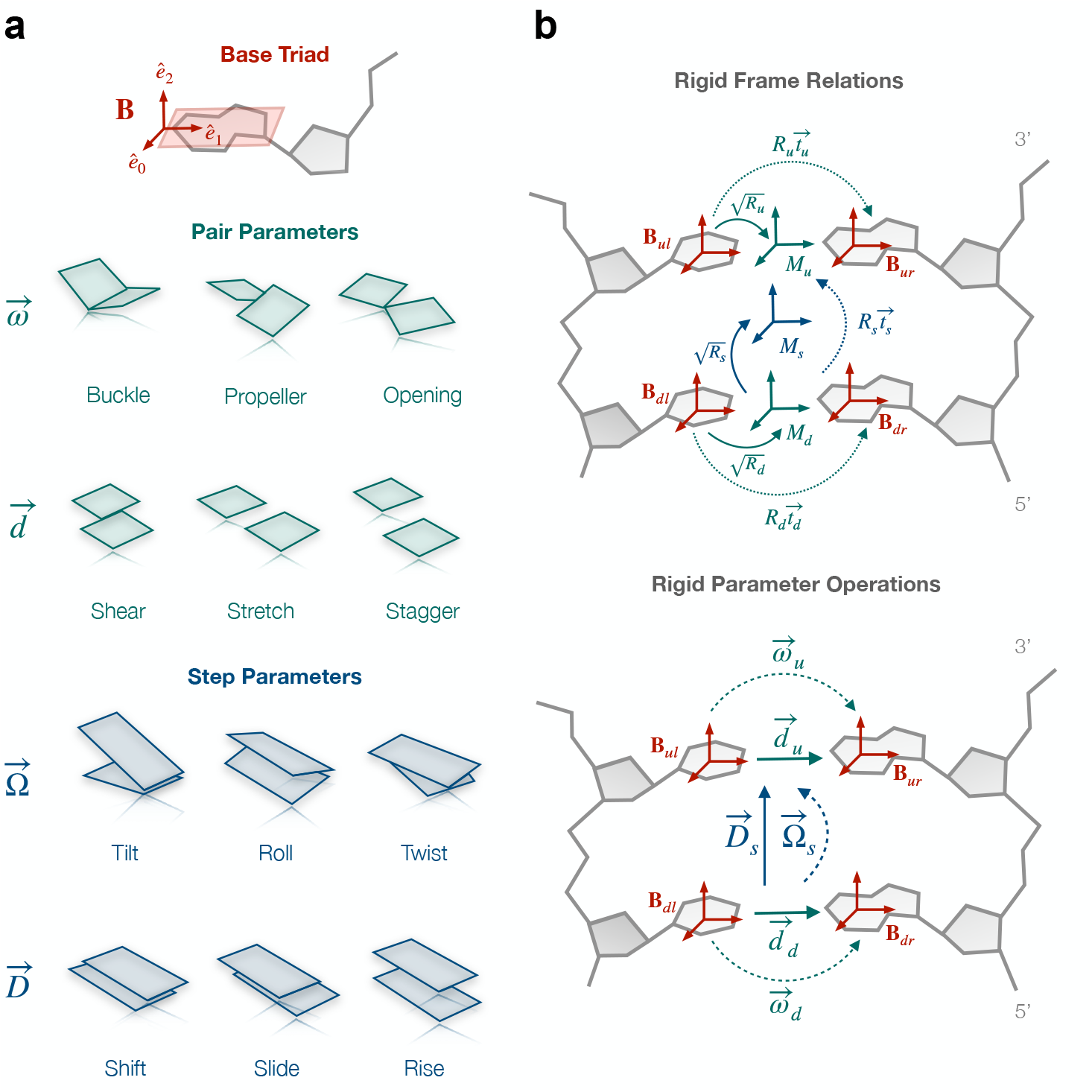
**a)** Top, an illustration of the unit vectors forming a base triad, also called the base reference frame. Intra-base pair (middle) and inter-base pair (bottom) parameters of a coarse-grain DNA model with the rotational parameters are on the top, translation parameters on the bottom. Not that the bases/base pairs are represented as cuboids. **b)** Top, diagram showing the relation between different rotations and reference triad. Bottom: from the reference frames = 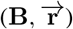 consisting of an orthonormal triad **B** and its origin 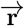, one calculates translational 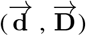 and rotational 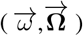 rigid base coordinates 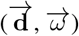 are intrabase coordinates, while 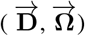 are interbase coordinates.

To parameterize translations and rotations connecting two complementary base frames on opposite strands of a DNA molecule, one uses the rigid base coordinates 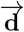 (a vector whose components are referred to as shear, stretch, and stagger in the DNA literature Wilma K Olson et al. 2001) and 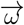 (buckle, propeller, and opening). The coordinates 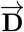 (shift, slide, and rise) and 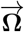 (tilt, roll, and twist) parameterize translations and rotations between two consecutive base pairs. See Figure 5-a for a visual representation of the rigid base parameters.

To obtain the translational and rotational parameters for each base and base pair, we start with the transformation between two opposing base frames (**B**_*ur*_ and **B**_*ul*_), characterized by a rotation matrix **R**_*u*_ and a translation vector **t**_*u*_:

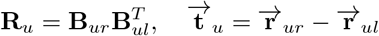

These transformations are expressed in a midframe **M**_*u*_, which is determined by rotating the vectors in **B**_*ul*_ by half the rotation angle of **R**_*u*_. Next, we compute the angle Θ (**R**) and the axis 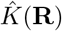 of rotation.

The angle Θ (**R**) and the axis 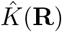 of rotation are given by:

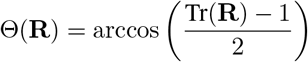

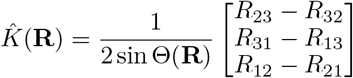

The rotational and translational parameters of the base pair are as follows:

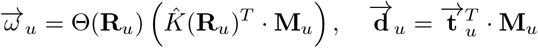

The process repeats for successive base pairs **B**_*dr*_ and **B**_*dl*_. After this, we can use the respective mid frames of the base pairs, **M**_*u*_ and **M**_*d*_, where the transformation between pairs involves a new rotation **R**_*s*_ and translation 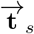:

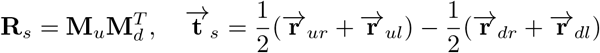

The midframe **M**_*s*_ for this step is defined analogously:

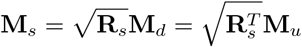

Finally, the transformations are expressed in this base step midframe, with the corresponding rotational and translational variables:

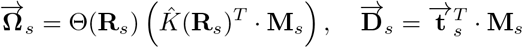

Figure 5-b shows a diagram of both the relations (top) and the operations (bottom) to obtain the rigid base parameters.

### 5.5 Calculation of Linking Number

Intuitively, the linking number Lk of two oriented closed curves 𝒞_1_ and 𝒞_2_ describes the frequency with which the curves wind around each other. Alternatively, it can be seen as the number of times one strand crosses any surface bounded by the other, formalized by Gauss’s linking integral Gauss, Gauss, and Wissenschaften zu Göttingen 1877:

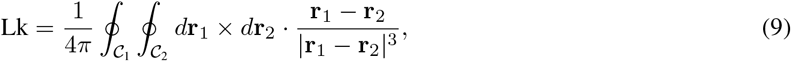

where **r**_1_ and **r**_2_ are the points along the two curves 𝒞_1_ and 𝒞_2_, respectively.

In a relaxed dsDNA molecule, the right-handed helical structure introduces one link per helical repeat (*n*_*h*_ = 10.5 bp or *h* = 3.57 nm for B-DNA). For a molecule of length *L* this results in a relaxed state linking number of *Lk*_0_ = *L/h*.

The sign of the linking depends on the directionality of the strand. For DNA, it is typically defined so that *Lk* is positive for right-handed helices and negative for left-handed ones. However, the linking number *Lk* may differ from *Lk*_0_ when the helix is over- or under-twisted, leading to an excess linking number Δ*Lk* = *Lk − Lk*_0_.

The linking number is a conserved topological property that always assumes integer values. However, according to the Călugăreanu-White-Fuller theorem Călugăreanu 1961; Fuller 1971; White 1968, the linking number of two parallel running space curves may be decomposed into two geometric quantities—twist and writhe—as

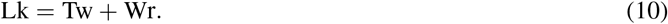

Twist indicates local winding of the two strands, whereas writhe indicates non-local coiling.

Twist, as a local property, involves a single integral over twist density, while writhe, a non-local property, requires a double integral akin to Gauss’ integral. For any closed, smooth curve𝒞, the writhe Wr is given by:

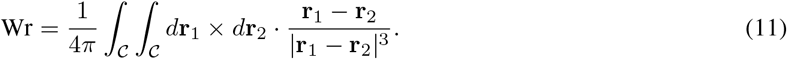

where **r**_1_ and **r**_2_ are points on the curve𝒞.

This integral spans the center line curve 𝒞 of the helix. Unlike linking, writhe is tied to a single space curve, reflecting the average signed crossings from all perspectives Klenin and Langowski 2000a and vanishing in fully planar curves or those with reflection symmetry.

In practice, we evaluate the linking number for a chain of straight segments, which naturally decomposes Gauss’s linking integral into a double sum of pairwise components. Contributions to the linking number stemming from such pairs of straight segments may be evaluated analytically; for details, see A. V. Vologodskii et al. 1979 and Method 1b in Ref. Klenin and Langowski 2000b. Similarly, for the numerical evaluation of Eq. 11 we rely on Method 1a of Ref. Klenin and Langowski 2000b.

### 5.6 Process of H-NS filament construction

H-NS (Histone-like nucleoid-structuring protein) is a bacterial DNA-binding protein that plays a crucial role in the organization of the nucleoid in gram-negative enterobacteria. The protein structures DNA by forming filaments along DNA duplexes, either by binding to two separate DNA duplexes or to adjacent sites on the same duplex Valk et al. 2017; Falconl et al. 1988; Williams and Rimsky 1997; Dame, Wyman, and Goosen 2000; Dorman 2004; Y. Liu et al. 2010. H-NS is made up of 137 amino acid residues, divided into two main domains: the oligomerization domain and the DNA binding domain (DBD). The oligomerization domain, which consists of residues 1-83, includes two key sites: the homodimerization site (s1) and the multimerization site (s2), which allow the formation of higher order structures Arold et al. 2010. The DBD, which spans residues 89-137, consists of an antiparallel *β*-sheet, an *α* helix, and a 3_10_ helix Shindo et al. 1995; Gordon et al. 2011. NMR experiments on the full H-NS protein indicated that the oligomerization domain and DBD function independently of each other, with a flexible linker connecting the two domains Dorman, Hinton, and Free 1999.

Given the absence of detailed structural data for H-NS filaments, the following protocol outlines a computational approach to construct H-NS filaments using available structural data and a protocol outlined by Van der Valk et al. Van Der Valk et al. 2015, see Figure 6. To construct a full-length H-NS monomer we used residues 2–83, from *S. typhimurium* (PDB 3NR7 Arold et al. 2010) and the NMR structure of the C-terminal domain, residues 91–137 (PDB 2L93 Gordon et al. 2011), while non-resolved sequences (residues 1 and 84–90) were modeled as a random coil. The structure of residues 2–83 of *S. typhimurium* H-NS also contained information on the dimerization sites. The coordinates of the DNA binding domain (DBD) of H-NS bound to the minor groove of a high-affinity 12-bp strand of dsDNA were obtained from earlier work Riccardi et al. 2019. Using these structures as templates, we combined 12 H-NS monomers into a multimer. To form the multimer, functional segments within the H-NS structure are identified: s1 (homodimerization domain), h3 (a helical region that potentially connects the s1 and s2 domains), s2 (multimerization domain), l2 (flexible linker region), and DBD (DNA binding domain). To form the multimer, we also included two structures as a building block where two s1 segments and two s2 segments are non-covalently bound. The segments are aligned using superposition techniques, where the root mean squared deviation (RMSD) of atom selections based on the overlapping residues between segments is minimized using singular value decomposition (SVD). Following alignment, the structures are joined to form a continuous H-NS filament. During this assembly process, the residue numbering is adjusted to maintain chain continuity. The initial conformations of the segments are selected based on short molecular dynamics (MD) simulations of the s1s1 and s2s2 system (see Figure 6) that ensure that the filament grows linearly, without intersecting with itself, causing overlap between the atomic positions. For computational details of the MD simulations, see section 5.7. The code to reproduce this protocol can be found on the documentation webpage.

**Figure 6:**
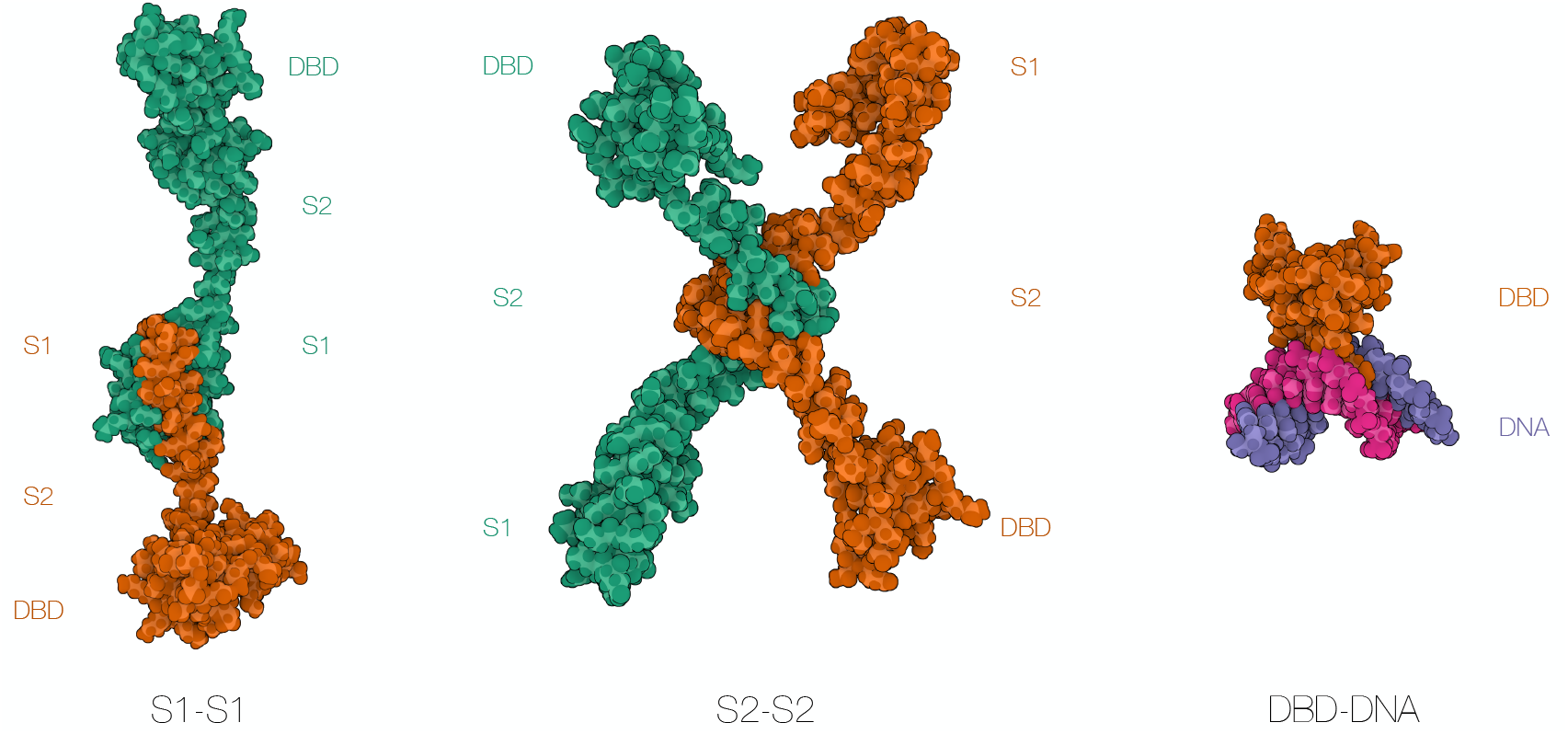
H-NS systems used to generate lateral protein filament: the H-NS homodimer (s1s1 system), the multimerization dimeric unit (s2s2 system), and the DNA binding domain (DBD) DNA complex. Coordinates of the DNA binding domain (DBD) of H-NS bound to the minor groove of a high-affinity 12-bp strand of dsDNA were obtained from earlier work Riccardi et al. 2019. Systems s1s1 and s2s2 were constructed using two monomers. The monomers were constructed based (PDB 3NR7 Arold et al. 2010 and PDB 2L93 Gordon et al. 2011). Molecular representations are visualized with Mol* Viewer Sehnal et al. 2021.

### 5.7 Molecular Simulations Methods of Nucleosome and H-NS dimers

Molecular dynamics (MD) simulations of the following systems have been performed using the protocol outlined below. The Nucleosome Core Particle with 147 bp DNA has been simulated for a duration of 250 ns for the analysis as shown in Section 2.4. For the construction of the H-NS filament as described in 5.6, the following MD simulations have been performed: For the s1s1 system, 15 runs of 62.5 ns starting from the same structural configuration were performed. Additionally, there are two longer runs of 100 ns each. These runs were initiated from a closed or open state characterized by the availability of the DNA binding domain (DBD) for DNA binding. For the s2s2 system, 12 simulations of 100 ns each were performed, all starting from the same initial structure.

The preparation of the systems for MD simulations consisted of placing the structures in a periodic dodecahedron box, with the box boundaries at least 1.0 nm from the system, followed by the addition of water molecules. To mimic experimental conditions Gordon et al. 2011 and neutralize the system, we added 50 mM NaCl by replacing water molecules with ions. The interactions between atoms are described by the force field AMBER14sb-parmbsc1 Maier et al. 2015; Ivani et al. 2016 in combination with the TIP3P water model Jorgensen et al. 1983. We selected this particular force field because it covers the topologies for both amino acids and nucleotides and provides good representations of the static and dynamic properties of DNA under a wide range of conditions Ivani et al. 2016. For non-bonded interactions, both van der Waals and electrostatic, we used a cut-off at 1.1 nm. Long-range electrostatic interactions were handled using the particle mesh Ewald method Cheatham et al. 1995; Essmann et al. 1995 with a grid spacing of 0.12 nm. To remove unfavorable interactions, we performed energy minimization using steepest descent. By applying position restraints on the heavy atoms of the protein and DNA with a force constant in each direction of 1000 kJ/mol nm^2^ and performing 0.1 ns of MD at a temperature of 298 K and a pressure of 1 bar, we relaxed the water and ions around the initial structures.

After preparation, we performed MD simulations varying initial conditions by assigning new random starting velocities drawn from the Maxwell-Boltzmann distribution at 298 K. All simulations were performed with GROMACS, version 2020.4 Van Der Spoel et al. 2005; Abraham et al. 2015 in a locally maintained cluster, with the leap-frog integration scheme and a time step of 2 fs, using LINCS Hess et al. 1997 to constrain the proteins and SETTLE Miyamoto and Kollman 1992 to constrain the water bonds. All simulations were performed in the isothermal-isobaric ensemble at a pressure of 1 bar, using the v-rescale thermostat Bussi, Donadio, and Parrinello 2007 and the isotropic Parrinello-Rahman barostat Parrinello and Rahman 1981; Nosé and Klein 1983.

